# Nanoporous Boron Nitride for High Efficient Water Desalination

**DOI:** 10.1101/500876

**Authors:** Zonglin Gu, Shengtang Liu, Xing Dai, Serena H. Chen, Zaixing Yang, Ruhong Zhou

**Author notes:** These authors contribute equally (Zonglin Gu, Shengtang Liu). Corresponding author, (Z.Y.); (R.Z.).

## Abstract

Membrane filtration processes for water desalination have been greatly improved thanks to rapid development of nanoporous 2-dimentional (2D) materials. Nanoporous graphene and molybdenum disulfide have proved to show promising properties for desalination. In this study, we detailly investigated the desalination performance of a different nanoporous 2D material, nanoporous boron nitride (BN), by Molecular Dynamics simulation. Our calculations demonstrated that nanoporous BN allows for rapid water permeability with effective salt rejection. The permeability is not only two orders of magnitude higher than existing commercial techniques but also much higher than nanoporous graphene and molybdenum disulfide membranes. We further showed that the pores with B-h edges or with N-h edges present different desalination efficiency. Compared to N-h pores, B-h pores have better desalination performance in term of higher water flux. To the best of our knowledge, nanoporous BN is the 2D material having the highest water permeability thus far while maintaining high salt rejection. Overall, our results shed light on the potential

## INTRODUCTION

Due to rapid consumption of freshwater, the governments of many countries are facing an increasing demand for freshwater resources. Aside from the small amount of available freshwater, about 97% of the world’s water is found in oceans and seas. These immense potential water resources have motivated research in desalination technologies to turn seawater into freshwater. Yet there are challenges in the development of desalination techniques.^1–2^ The pivotal factors are high capital costs and low efficiency which, on the other hand, motivate advances in this field.^3^ Reverse Osmosis (RO) is the most popular desalination method. The process includes (i) placing a RO membrane at the interface between seawater and freshwater, and (ii) applying pressure at the seawater side to facilitate the flow of water molecules through the RO membrane to the freshwater side while leaving salt ions behind. Despite the wide use of the technique, the overall desalination performance of existing commercial RO membranes is still under satisfaction due to their slow water transport. Thus, developing high-efficiency membrane filter is one of the critical solutions to improve the efficiency of RO desalination.

Thanks to rapid development of nanoporous 2-dimentional (2D) materials (through either ion irradiation or chemical treatment)^4–5^, membrane filter preparation for water desalination has been greatly improved. The materials containing nanopores with diameters ranging from several angstroms to a few nanometers have been widely used in molecular sieves.^6–8^ Some nanopores with pore sizes smaller than the size of hydrated ions are suitable as membrane filters as they retain the ions but allow passage of water.^9–10^ In addition to pore size, membrane thickness proved to be negatively correlated with water flux. A typical example is single-layered nanoporous graphene (including chemically modified nanopores), which proved to have several orders of magnitude higher flux rates than commercial RO membranes.^11^ Interestingly, Ghoufi *et al.* recently found that in pure water under the same aperture (~ 7 Å), BN nanoporous with an approximately circular channel possesses an even higher water permeability than graphene nanosheets, due to the smaller surface tension of water on BN nanosheet.^12^ Another example of RO membranes is three-atom layered nanoporous molybdenum disulfide (MoS_2_), which also has higher water flux than commercial RO membranes but just a little lower water flux than nanoporous graphene.^13^ Owing to the three-atom layers (S-Mo-S), MoS_2_ nanopores can also be used as molecular switches, which provide “open” and “closed” states, for water transport by applying proper lateral strain in desalination.^14^

Herein, we investigated a novel triangular nanoporous 2D material, nanoporous boron nitride (BN), for rapid and effective desalination using Molecular Dynamics (MD) simulation. Compared to commercial desalination techniques, nanoporous BN shows higher efficiency of water permeation and salt rejection. Our study provides a new material for RO in future water desalination.

## RESULTS

The schematic view of the simulation setup is depicted in Fig. 1a in which four major components of the system are labeled. We calculated the water permeability through two different types of BN nanopores, N-h (Fig. 1b) and B-h (Fig. 1c) pores (definitions of pore types can be found in METHODS section). The pores were designed in triangle shape based on the unique feature of BN nanosheet recently releveled by transmission electron microscopy (TEM) experiments, where triangular nanopores can be fabricated and regulated while retaining their intrinsic triangular shape.^15–18^ The number of filtered water molecules across the pores was monitored over simulation time by calculating the amount of increase in the number of water molecules at the fresh water side (Fig. 2a). The results demonstrate a linear relationship between the number of filtered water molecules and simulation time. Moreover, the filtration rate increases as the external pressure applied to the piston increases. It is worth to note that the number of filtered water molecules through B-h pores is higher than that of N-h pores at the same pressure, and the difference increases as the pressure increases (Fig. 2b). Besides, with the same pore size, B-h pores have noticeable higher water permeability than N-h pores (Fig. 2c; more below). Whereas, when the pore size narrowed down to N-h-4 pore (with pore area of ~23.1 Å^2^), the flow of water starts to shut off.

**Figure 1.**
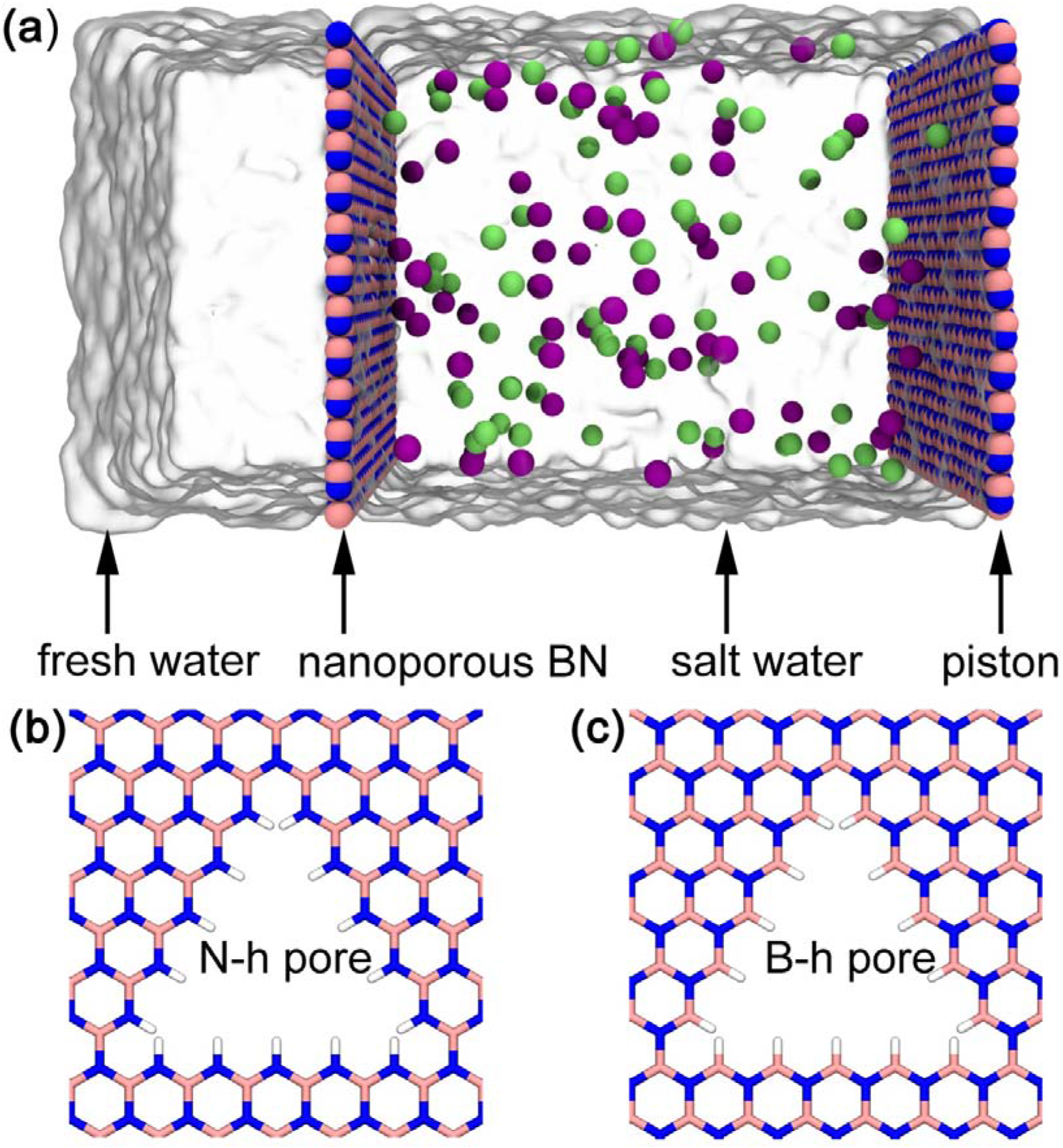
(a) Schematic view of the simulation setup. The system consists of a piston (BN nanoplate), salt water (including sodium and chloride ions), a separation membrane (nanoporous BN nanoplate), and fresh water. The boron and nitrogen atoms of the two BN nanoplates are represented by pink and blue spheres, respectively. Sodium ions are colored in lime and chloride ions are in mauve. (b and c) Two types of nanopores. (b) A nanopore with nitrogen and hydrogen edges (N-h pore). (c) A nanopore with boron and hydrogen edges (B-h pore).

**Figure 2.**
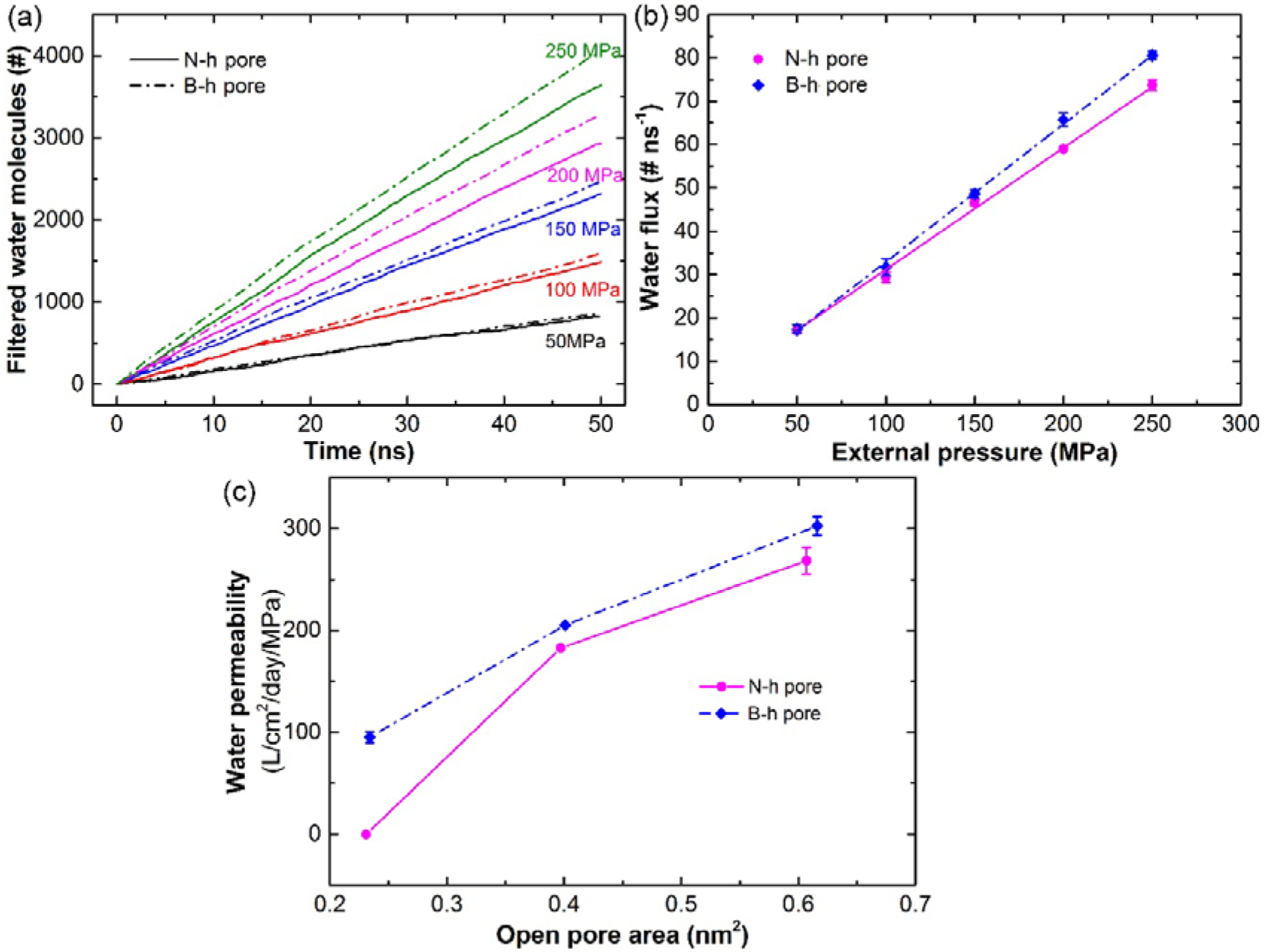
(a) Number of filtered water molecules through the nanoporous BN nanoplate with N-h-5 pores (solid lines) and with B-h-5 pores (dot-and-dash lines) at piston pressure ranging from 50 to 250 MPa. (b) Pressure dependence of water flux through the two types of BN nanopores (B-h-5 and N-h-5 pores). (c) Computed water permeability of N-h and B-h pores with three different pore sizes.

In addition to water permeability, we computed salt rejection of different pore types and pore sizes based on the salinity of the permeate solution at t = t_1/2_ relative to the initial salinity,^11^ where t_1/2_ is defined as the time point when half of the water molecules has flowed to the fresh water side. The results are shown in Fig 3. Compared to other nanopores, B-h-4 pore has the most efficient salt rejection, which reaches 100%. On the other hand, N-h-4 pore has zero water permeability at the pressure values applied in this study, and therefore, we do not show its salt rejection. The salt rejections of B-h-5 pore (mean value of 93.5%) and N-h-5 pore (mean value of 94.7%) are comparable, while those of B-h-6 pore and N-h-6 pore have much lower ratios. These results agree with previous observations that salt rejection is lower as the pore size becomes bigger.^11, 13^ Overall, our results indicate that both water permeability and salt rejection of nanoporous BN are specific to its pore sizes.

**Figure 3.**
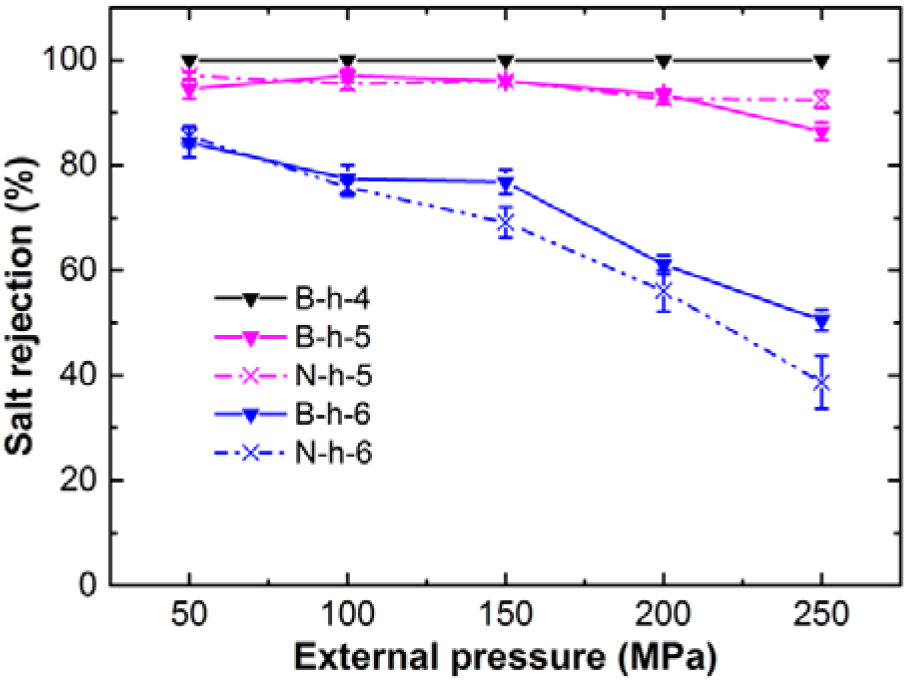
Pressure-dependent average salt rejection of the two types of BN nanopores with three different pore sizes. N-h-4 pore is not shown because it has zero water permeability at the pressure values applied.

**Figure 4.**
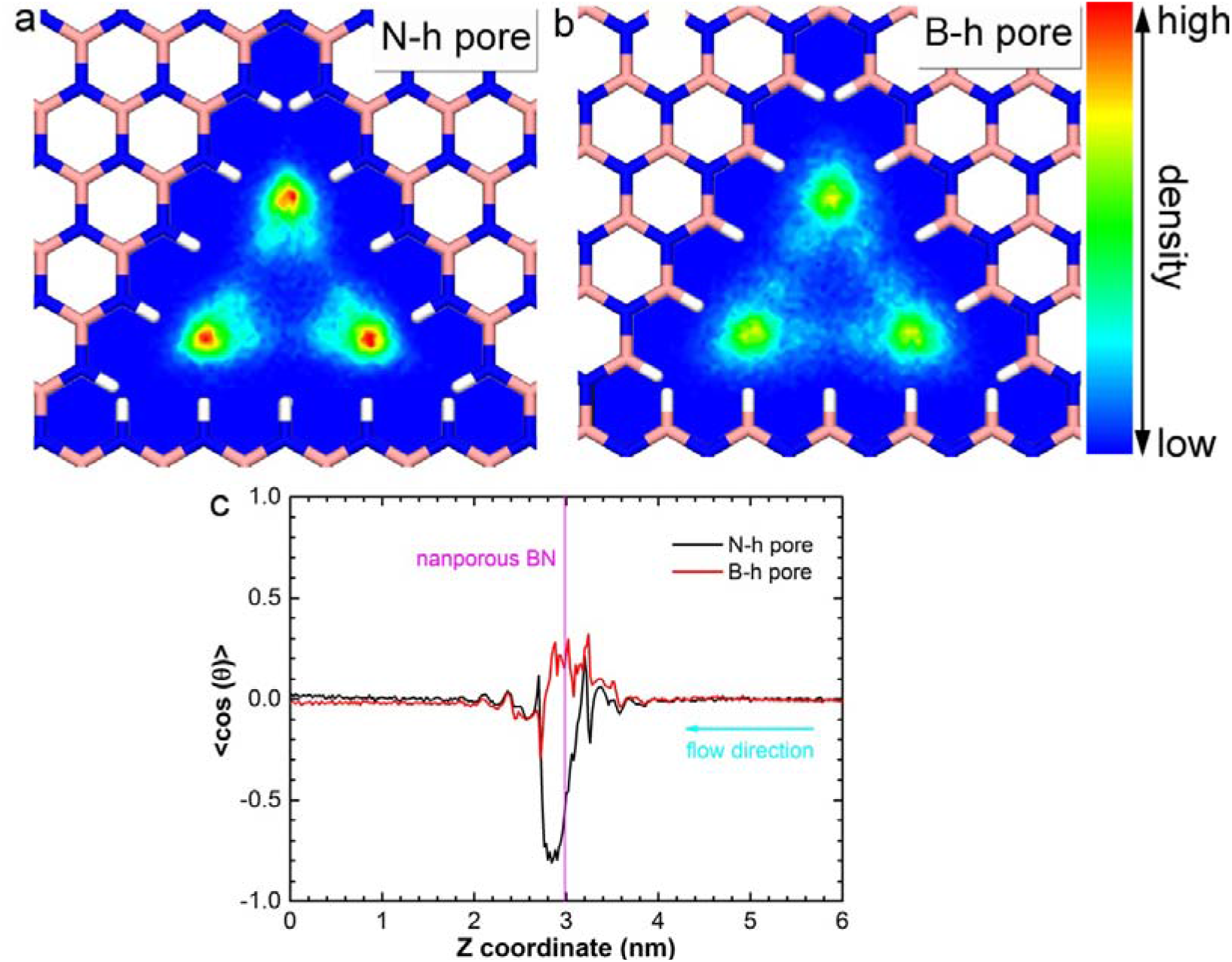
Water density maps inside a N-h-5 pore (a) and a B-h-5 pore (b). No water molecule was found in the blue regions, while the probability of finding a water molecule was the highest in the red regions. (c) Average orientation, <cos(θ)>, of water molecules with respect to the normal direction (the z coordinate) of nanoporous BN nanoplate, which is indicated by the perpendicular line in magenta. The cyan arrow points toward the direction of water flow.

The orientation of water molecules near the pores plays a critical role in both water permeability and salt rejection of nanoporous BN. The water density maps inside the two types of the pores are depicted in Fig. 4a and 4b. Water molecules in the N-h pore are highly localized at the three corners of the pore, while water molecules in the B-h pore distribute relatively evenly and the density values at the three corners are not as high as those in the N-h pore. However, the mean water density inside the B-h pore is ~1.945 water molecules per nm^3^, which is slightly higher than that in the N-h pore (~1.824 water molecules per nm^3^), implying that the B-h pore can accommodate more waters to traversing the filter membrane than N-h pore at similar pore area which might help decipher the higher water permeability of B-h pore than N-h pore. Besides, the average orientation, <cos(θ)>, of water molecules with respect to the normal direction (the z coordinate) of the nanoporous BN nanoplate is shown in Fig. 4c. Here *θ* represents the angle between the dipole moment of water and the z-axis of the simulation box. The values change from positive to negative along the flow direction, suggesting that water molecules rotate when passing through the membrane filter. The rotation angle of the water molecules in the N-h pore is very large (<cos(θ)> changes from 0.212 to −0.809, i.e., from 77.8° to 144.0°). On the other hand, the water molecules do not rotate much when crossing the B-h pore (<cos(θ)> changes from 0.281 to −0.290 (i.e., from 73.7° to 106.9°). Instead, the water molecules rotate after passing the pore at the position of 0.1 nm from the BN nanoplate. The smaller rotation angle offers a smoother entropic landscape for water molecules to traverse and thus results in a faster water flow.

We also compared the desalination performance between nanoporous BN and existing desalination techniques based on their water permeability and salt rejection (Fig. 5). The B-h-4 pore showed 100% salt rejection and high water passage. The water permeability is 95.3 L per cm^2^ day MPa, which is not only two orders of magnitude higher than that of commercial technologies but much higher than that observed with nanoporous graphene or MoS_2_ membrane.^11, 13^ Additionally, the N-h-5 and B-h-5 pores present more efficient permeability than the B-h-4 pore while their salt rejections are slightly lower (94.7% and 93.5%, respectively). To our best knowledge, the nanoporous BN is the best RO membrane so far compared to other materials.

**Figure 5.**
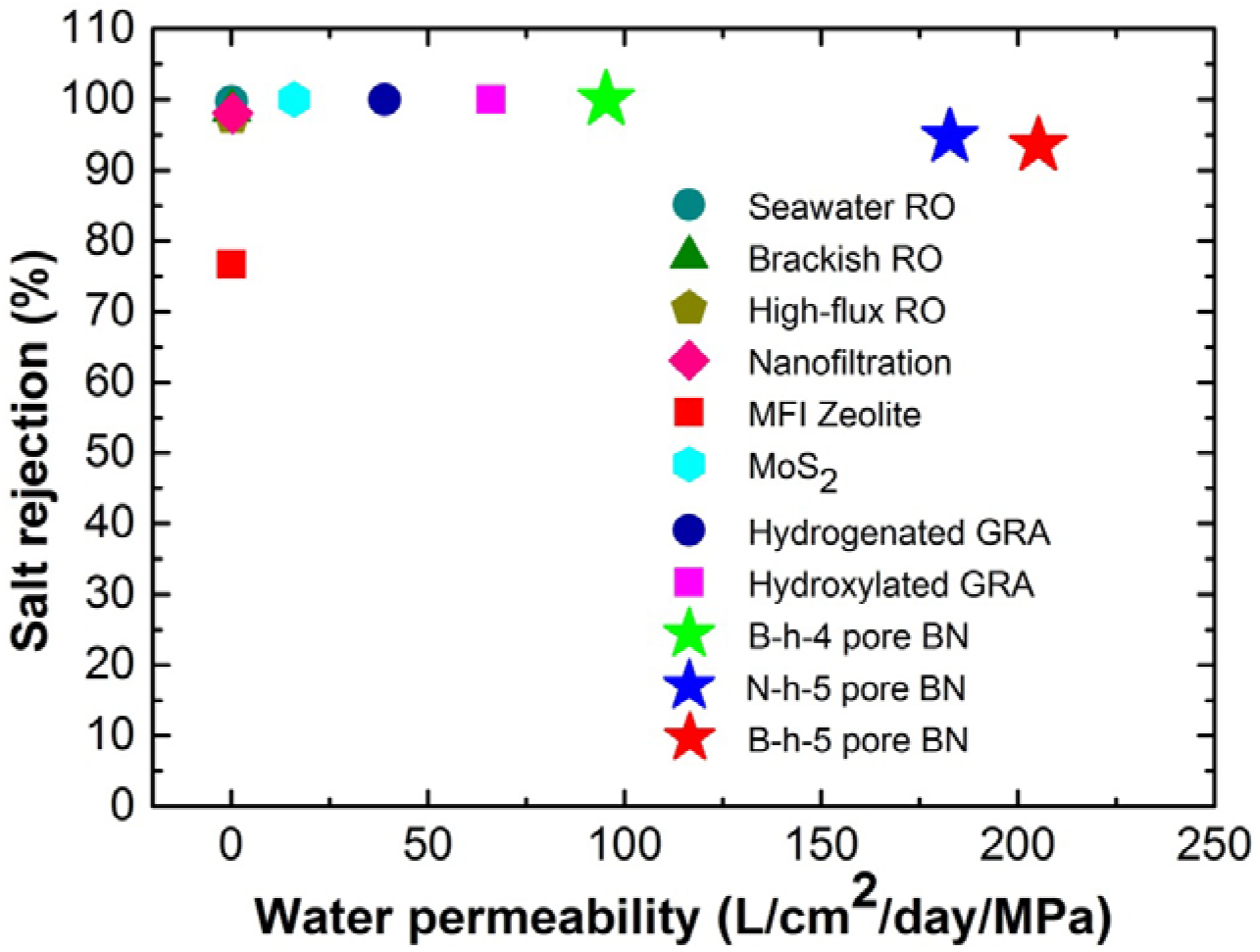
Desalination performance of nanoporous BN and existing desalination techniques. The data of commercial RO and MFI zeolites, MoS_2_ and hydrogenated/hydroxylated graphene (GRA) were adapted from Pendergast et al,^19^ Heiranian et al^13^ and Cohen-Tanugi et al,^11^ respectively.

## CONCLUSION

We demonstrated that nanoporous boron nitride has outstanding desalination capacity with rapid water permeability and high salt rejection. B-h pores allow for larger water flux than N-h pores as the water molecules undergo dipole rotation inside N-h pores which slows down the water flow. Compared to existing desalination techniques, nanoporous boron nitride has overall better desalination performance. Our findings have shed light on a new candidate for future desalination membrane filter.

## METHODS

There are two types of system in our simulations: the nanopores with N-h edges and those with B-h edges. Each system consists of a piston (a BN sheet), salt water, a filter (a nanoporous BN sheet) and pure water (as shown in Fig. 1a). For each of the two pore types, three different pore sizes were considered –M-h-4, M-h-5 and M-h-6 (M = N, B). The M-h-4 pore has four M-h pairs at every side of the triangular pore. Similarly, the M-h-5 and M-h-6 pores have five and six M-h pairs at each side of the pore, respectively. The open pore areas for B-h-4 pore, B-h-5 pore, B-h-6 pore, N-h-4 pore, N-h-5 pore, and N-h-6 pore are about 0.234 nm^2^, 0.401 nm^2^, 0.611 nm^2^, 0.231 nm^2^, 0.397 nm^2^, and 0.607 nm^2^, respectively. Following similar protocols in our previous studies^20–31^, these nanopores are solvated in water boxes with dimensions of the system at 5.00 nm ×5.21 nm ×10.00 nm in x, y and z directions, respectively. The box contains a total of ~8300 water molecules and 0.599 M NaCl in salt water (to mimic the salinity of seawater).

Molecular Dynamics simulations were performed using the GROMACS software package (version 4.6.6)^32^. The VMD software^33^ was used to analyze and visualize the simulation results. CHARMM27 force field^34–35^ and TIP3P^36^ water model were used for the Na^+^/Cl^−^ and water molecules, respectively. The force field parameters of BN were obtained from a previous study.^12^ Temperature was fixed at 300 K using v-rescale thermostat^37^. Periodic boundary conditions were applied in all directions. The long-range electrostatic interactions were calculated by the PME method^38–39^, and the van der Waals (vdW) interactions were calculated with a cutoff distance of 1.2 nm. All solute bonds were kept constant at their equilibrium values with the LINCS algorithm^40^, and water geometry was constrained using the SETTLE algorithm^41^. During the production runs, a time step of 2.0 fs was used. Three independent trajectories were generated for each system. All simulations were performed for over 50 ns to ensure the condition of salt rejection where half of the water molecules in the salt water side has across the nanoporous BN nanoplate.

## REFERENCES

1. Elimelech M.; Phillip, W. A., The Future of Seawater Desalination: Energy, Technology, and the Environment. Science 2011, 333 (6043), 712-717.

2. Shannon M. A.; Bohn, P. W.; Elimelech, M.; Georgiadis, J. G.; Marinas, B. J.; Mayes, A. M., Science and Technology for Water Purification in the Coming Decades. Nature 2008, 452 (7185), 301–310.

3. Khawaji A. D.; Kutubkhanah, I. K.; Wie, J.-M., Advances in Seawater Desalination Technologies. Desalination 2008, 221 (1-3), 47–69.

4. Surwade S. P.; Smirnov, S. N.; Vlassiouk, I. V.; Unocic, R. R.; Veith, G. M.; Dai, S.; Mahurin, S. M., Water Desalination Using Nanoporous Single-Layer Graphene. Nat. Nanotechnol. 2015, 10 (5), 459–464.

5. Merchant C. A.; Healy, K.; Wanunu, M.; Ray, V.; Peterman, N.; Bartel, J.; Fischbein, M. D.; Venta, K.; Luo, Z.; Johnson, A. T. C., et al., DNA Translocation through Graphene Nanopores. Nano Lett. 2010, 10 (8), 2915–2921.

6. Suk M. E.; Aluru, N. R., Water Transport through Ultrathin Graphene. J. Phys. Chem. Lett. 2010,1 (10), 1590–1594.

7. O’Hern S. C.; Stewart, C. A.; Boutilier, M. S. H.; Idrobo, J.-C.; Bhaviripudi, S.; Das, S. K.; Kong, J.; Laoui, T.; Atieh, M.; Karnik, R., Selective Molecular Transport through Intrinsic Defects in a Single Layer of Cvd Graphene, Acs Nano 2012, 6 (11), 10130–10138.

8. Zhao Y.; Xie, Y.; Liu, Z.; Wang, X.; Chai, Y.; Yan, F., Two-Dimensional Material Membranes: An Emerging Platform for Controllable Mass Transport Applications. Small 2014,10 (22), 4521–4542.

9. Zhang D.; Yan, T.; Shi, L.; Peng, Z.; Wen, X.; Zhang, J., Enhanced Capacitive Deionization Performance of Graphene/Carbon Nanotube Composites. J. Mater. Chem. 2012, 22 (29), 14696–14704.

10. Celebi K.; Buchheim, J.; Wyss, R. M.; Droudian, A.; Gasser, P.; Shorubalko, I.; Kye, J.-I.; Lee, C.; Park, H. G., Ultimate Permeation across Atomically Thin Porous Graphene. Science 2014, 344 (6181), 289–292.

11. Cohen-Tanugi, D.; Grossman, J. C., Water Desalination across Nanoporous Graphene. Nano Lett. 2012,12 (7), 3602–3608.

12. Gamier L.; Szymczyk, A.; Malfreyt, P.; Ghoufi, A., Physics Behind Water Transport through Nanoporous Boron Nitride and Graphene. J. Phys. Chem. Lett. 2016, 7 (17), 3371–3376.

13. Heiranian M.; Farimani, A. B.; Aluru, N. R., Water Desalination with a Single-Layer Mos2 Nanopore. Nat. Commun. 2015, 6, 8616.

14. Li W.; Yang, Y.; Weber, J. K.; Zhang, G.; Zhou, R., Tunable, Strain-Controlled Nanoporous Mos2 Filter for Water Desalination. Acs Nano 2016,10 (2), 1829–1835.

15. Meyer J. C.; Chuvilin, A.; Algara-Siller, G.; Biskupek, J.; Kaiser, U., Selective Sputtering and Atomic Resolution Imaging of Atomically Thin Boron Nitride Membranes. Nano Lett. 2009, 9 (7), 2683–2689.

16. Jin C.; Lin, F.; Suenaga, K.; Iijima, S., Fabrication of a Freestanding Boron Nitride Single Layer and Its Defect Assignments. Phys. Rev. Lett. 2009,102 (19), 195505.

17. Alem N.; Erni, R.; Kisielowski, C.; Rossell, M. D.; Gannett, W.; Zettl, A., Atomically Thin Hexagonal Boron Nitride Probed by Ultrahigh-Resolution Transmission Electron Microscopy. Phys. Rev. B 2009, 80 (15), 155425.

18. Liu K.; Lihter, M.; Sarathy, A.; Caneva, S.; Qiu, H.; Deiana, D.; Tileli, V.; Alexander, D. T. L.; Hofmann, S.; Dumcenco, D., et al., Geometrical Effect in 2d Nanopores. Nano Lett. 2017, 17 (7), 4223–4230.

19. Pendergast M. M.; Hoek, E. M. V., A Review of Water Treatment Membrane Nanotechnologies. Energy Environ. Sci. 2011, 4 (6), 1946–1971.

20. Stirnemann G.; Kang, S. G.; Zhou, R.; Berne, B. J., How Force Unfolding Differs from Chemical Denaturation. Proc. Natl. Acad. Sci. U. S. A. 2014, 111 (9), 3413–8.

21. Das P.; Li, J.; Royyuru, A. K.; Zhou, R., Free Energy Simulations Reveal a Double Mutant Avian H5n1 Virus Hemagglutinin with Altered Receptor Binding Specificity. Journal of computational chemistry 2009, 30 (11), 1654–1663.

22. Xia Z.; Clark, P.; Huynh, T.; Loher, P.; Zhao, Y.; Chen, H.-W.; Rigoutsos, I.; Zhou, R., Molecular Dynamics Simulations of Ago Silencing Complexes Reveal a Large Repertoire of Admissible ‘Seed-Less’ Targets. Scientific reports 2012, 2, 569.

23. Xiu P.; Yang, Z.; Zhou, B.; Das, P.; Fang, H.; Zhou, R., Urea-Induced Drying of Hydrophobic Nanotubes: Comparison of Different Urea Models. The Journal of Physical Chemistry B 2011,115 (12), 2988–2994.

24. Li J.; Liu, T.; Li, X.; Ye, L.; Chen, H.; Fang, H.; Wu, Z.; Zhou, R., Hydration and Dewetting near Graphite-Ch(3) and Graphite-Cooh Plates. J. Phys. Chem. B 2005,109 (28), 13639–48.

25. Zhou R., Exploring the Protein Folding Free Energy Landscape: Coupling Replica Exchange Method with P3me/Respa Algorithm. J. Mol. Graph. Model. 2004, 22 (5), 451–63.

26. Zhu L.; Sheng, D.; Xu, C.; Dai, X.; Silver, M. A.; Li, J.; Li, P.; Wang, Y.; Wang, Y.; Chen, L., et al., Identifying the Recognition Site for Selective Trapping of (99)Tco4(-) in a Hydrolytically Stable and Radiation Resistant Cationic Metal-Organic Framework. J. Am. Chem. Soc. 2017, 139 (42), 14873–14876.

27. Zhou R.; Gao, H., Cytotoxicity of Graphene: Recent Advances and Future Perspective. Wiley Interdiscip Rev Nanomed Nanobiotechnol 2014, 6 (5), 452–74.

28. Fitch B. G.; Rayshubskiy, A.; Eleftheriou, M.; J., C. T.; Giampaga, M.; Zhestkov, Y.; Pitman, M. C.; Suits, F.; Grossfield, A.; Pitera, J., et al., Blue Matter: Strong Scaling of Molecular Dynamics on Blue Gene/L. Springe rBerlin Heidelberg: 2006.

29. Li X.; Li, J.; Eleftheriou, M.; Zhou, R., Hydration and Dewetting near Fluorinated Superhydrophobic Plates. J. Am. Chem. Soc. 2006,128 (38), 12439–47.

30. Das P.; Zhou, R., Urea-Induced Drying of Carbon Nanotubes Suggests Existence of a Dry Globule-Like Transient State During Chemical Denaturation of Proteins. J. Phys. Chem. B 2010,114 (16), 5427–30.

31. Kaminski G. A.; Friesner, R. A.; Zhou, R., A Computationally Inexpensive Modification of the Point Dipole Electrostatic Polarization Model for Molecular Simulations. J. Comput. Chem. 2003, 24 (3), 267–76.

32. Hess B.; Kutzner, C.; van der Spoel, D.; Lindahl, E., Gromacs 4: Algorithms for Highly Efficient, Load-Balanced, and Scalable Molecular Simulation. J. Chem. Theory Comput. 2008, 4 (3), 435–447.

33. Humphrey W.; Dalke, A.; Schulten, K., Vmd: Visual Molecular Dynamics. J. Mol. Graph. Model. 1996,14(1), 33–38.

34. Mackerell A. D.; Feig, M.; Brooks, C. L., Extending the Treatment of Backbone Energetics in Protein Force Fields: Limitations of Gas-Phase Quantum Mechanics in Reproducing Protein Conformational Distributions in Molecular Dynamics Simulations. J. Comput. Chem. 2004, 25 (11), 1400–1415.

35. MacKerell A. D.; Bashford, D.; Bellott, M.; Dunbrack, R. L.; Evanseck, J. D.; Field, M. J.; Fischer, S.; Gao, J.; Guo, H.; Ha, S., et al., All-Atom Empirical Potential for Molecular Modeling and Dynamics Studies of Proteins. J. Phys. Chem. B 1998,102 (18), 3586–3616.

36. Jorgensen W. L.; Chandrasekhar, J.; Madura, J. D.; Impey, R. W.; Klein, M. L., Comparison of Simple Potential Functions for Simulating Liquid Water. J. Chem. Phys. 1983, 79 (2), 926–935.

37. Bussi G.; Donadio, D.; Parrinello, M., Canonical Sampling through Velocity Rescaling. J. Chem. Phys. 2007,126(1), 014101.

38. Essmann U.; Perera, L.; Berkowitz, M. L.; Darden, T.; Lee, H.; Pedersen, L. G., A Smooth Particle Mesh Ewald Method. J. Chem. Phys. 1995,103 (19), 8577–8593.

39. Darden T.; York, D.; Pedersen, L., Particle Mesh Ewald – an N.Log(N) Method for Ewald Sums in Large Systems. J. Chem. Phys. 1993, 98 (12), 10089–10092.

40. Hess B.; Bekker, H.; Berendsen, H. J. C.; Fraaije, J., Lincs: A Linear Constraint Solver for Molecular Simulations. J. Comput. Chem. 1997,18 (12), 1463–1472.

41. Miyamoto S.; Kollman, P. A., Settle – an Analytical Version of the Shake and Rattle Algorithm for Rigid Water Models. J. Comput. Chem. 1992,13 (8), 952–962.

